# The small and large ribosomal subunits depend on each other for stability and accumulation

**DOI:** 10.1101/384362

**Authors:** Brian Gregory, Nusrat Rahman, Ananth Bommakanti, Md Shamsuzzaman, Mamata Thapa, Alana Lescure, Janice M. Zengel, Lasse Lindahl

## Abstract

The 1:1 balance between the numbers of large and small ribosomal subunits can be disturbed by mutations that inhibit the assembly of one of the subunits. We have investigated if the cell has mechanisms to counteract an imbalance of subunits. We show that accumulation of 40S subunits stops after abrogating 60S assembly. In contrast, cessation of the 40S pathways does not prevent 60S accumulation, but does, however, lead to fragmentation of the 25S rRNA in 60S subunits. Even though 40S subunits do not accumulate in the absence of 60S assembly, 40S assembly continues, indicating that excess 40S is degraded after assembly. Thus, the cell expends substantial resources in a futile attempt to increase the number of functional 80S ribosomes. We propose that a mechanism for preventing the accumulation of 40S subunits in excess over 60S subunits may have evolved to prevent mRNAs from being tied up in 40S-mRNA translation initiation complexes that cannot be turned into productive translation complexes because of the deficit of 60S subunits.

## Introduction

Ribosomes consist of two different subunits, both of which are required for translation. The small subunit (“40S” in eukaryotes) decodes the genetic message and the large subunit (“60S” in eukaryotes) catalyzes peptide bond formation. The biogenesis of the eukaryotic 40S and 60S ribosomal subunits is a complex process that is investigated most thoroughly in *Saccharomyces cerevisiae* (yeast), but essential features are largely conserved from yeast to humans (Tafforeau et al. 2013). The process begins in the nucleolus, continues in the nucleoplasm, and is completed in the cytoplasm (for in depth reviews, see (Woolford and Baserga 2013; Kressler et al. 2017)). The rRNA of the 40S subunit (18S) and two of the three rRNA components of the 60S subunit (5.8S and 25S) are all transcribed as a single precursor RNA (35S) by RNA Polymerase I, while the third rRNA component of the 60S (5S) is polymerized separately by RNA Polymerase III. During its transcription, or soon after, the long pre-rRNA is cleaved into two parts, each destined for one of the two ribosomal subunits (Osheim et al. 2004; Kos and Tollervey 2010; Talkish et al. 2016). After this split, the pre-40S and pre-60S ribonucleoprotein particles assemble along separate pathways (Woolford and Baserga 2013; Kressler et al. 2017).

The precursor rRNA transcripts are processed into the mature rRNA moieties concurrent with the incorporation of r-proteins into the pre-ribosomal subunits.

Ribosomal proteins are added to the precursor ribosomes in hierarchical waves, i.e., initially only a subset of r-proteins binds to the earliest precursor particles, while binding of proteins in subsequent waves depend on proteins assembled in the previous wave. Depending on their place in this hierarchy, r-proteins are referred to as “early”, “middle” and “late” binding proteins (Shajani et al. 2011; Gamalinda et al. 2014; de la Cruz et al. 2015; Zhang et al. 2016), although reduced growth rates and stressful conditions might perturb this order to some extent (Talkish et al. 2016). Besides the actual ribosomal components, the biogenesis of ribosomes requires over 250 protein and RNA factors that coordinate assembly, modify ribosomal components, and control ribosomal morphopoiesis (Woolford and Baserga 2013; de la Cruz et al. 2015; Turowski and Tollervey 2015; Kressler et al. 2017).

The cotranscription of rRNA for both subunits normally helps to ensure production of approximately equal numbers of the subunits, which is important for efficient protein translation and cost-effective use of the enormous amount of resources dedicated to ribosome production. However, the balance of ribosomal subunits can be upset by mutations in genes encoding ribosomal components and assembly factors that distort the assembly of one or the other subunit, a source of many human diseases (Mattijssen et al. 2010; Narla and Ebert 2010; Danilova and Gazda 2015; Farley and Baserga 2016; Bustelo and Dosil 2018). It is not known if mechanisms have evolved to ameliorate subunit imbalance. In this study, we used yeast to investigate how abrogating assembly of one subunit affects the accumulation of the other. Our results show that obstructing 60S subunit assembly inhibits accumulation of 40S subunits due to post-assembly turnover. On the other hand, inhibiting 40S assembly does not prevent 60S subunit accumulation, although, interestingly, it does result in fragmentation of the 25S rRNA and structurally changed 60S subunits.

## Results

### Experimental set-up

We used strains that allow repression of the synthesis of a single ribosomal protein or assembly factor. In each strain, referred to as P_gal_-protein x, the only gene for protein x is transcribed from the *GAL1/10* promoter. Thus, protein x is synthesized in galactose medium, but not in glucose medium. Ribosomal proteins are referred to by the recently developed universal nomenclature (Ban et al. 2014), but classic yeast names are also shown on figures.

### Abrogating assembly of the 60S subunit inhibits accumulation of the 40S particles

We first disrupted the 60S assembly pathway at different points by repressing genes for early (uL4), middle (uL22), or late binding proteins (uL18 and eL43) or 60S assembly factors Rpf2 and Rrs1. In all cases, sucrose gradients analysis showed that the 60S peak was markedly reduced, but the 40S peak was increased relative to other ribosomal peaks (Figure 1, Supplementary Figure 1A-B), suggesting that 40S subunits were still produced, even though 60S assembly was blocked. Furthermore, all ribosomal peaks became smaller, although all gradients were loaded with the same amount of A^260^ material, showing that the ribosome content was diluted, as expected because ribosomes made prior to the repression of ribosomal genes continue to synthesize bulk protein. The ratio between the ribosomal peaks and A^260^ material remaining to the top of the gradient also decreased (Supplementary Figure 1H-I). Importantly, the same result was seen irrespective of which 60S r-protein or assembly factor gene was repressed, showing that the effect does not depend on which step in the 60S assembly pathway is disrupted.

**Figure 1.**
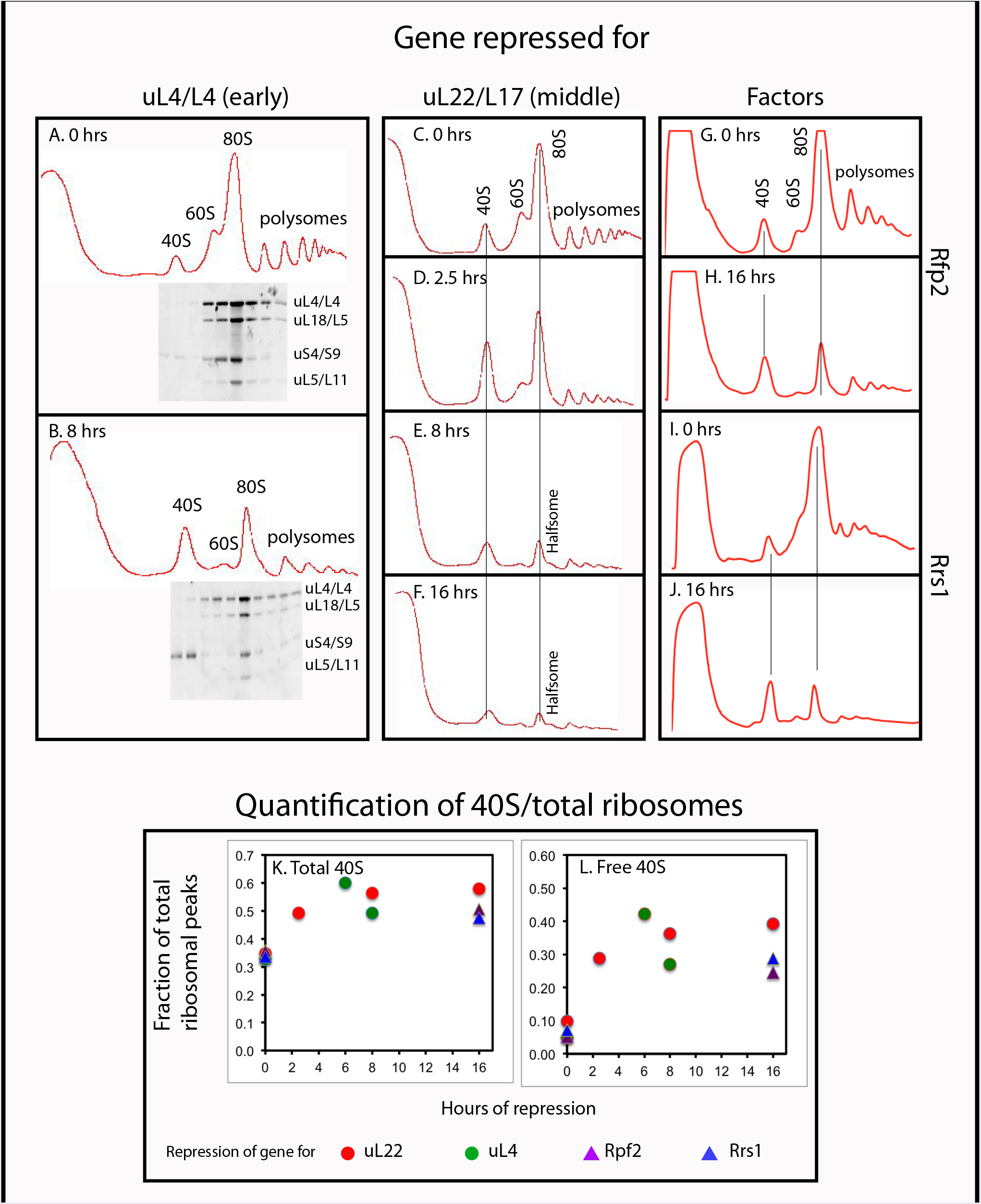
Sucrose gradient analysis of ribosomes after repressing the genes for 60S r-proteins uL4 and uL22 and 60S assembly factors Rpf2 and Rrs1. Cultures were grown in YEP galactose medium and shifted to YEP glucose medium for the indicated lengths of time before harvest. A-B. P_gal_-uL4 0 and 8 hours, respectively. C-F P_gal_-uL22 0, 2.5, 8, and 16 hours, respectively. G-H. P_gal_-Rpf2 0 and 16 hours, respectively. I-J. P_gal_-Rrs1 0 and 16 hours, respectively. For A and B, r-proteins uS4, uL4, uL5, and uL18 in the ribosomal fractions were analyzed by Western blot. K-L. Enumeration of 40S subunits. The area under each ribosomal peek was quantified and normalized to the area under all ribosomal peaks. K. The fraction of ribosomal mass in 40S subunits was calculated as the fraction found in the free 40S peak plus 1/3 of the fraction in 80S and polysomes. L. The fraction of ribosomal mass in free 40S subunits. Red and green circles refer to repression of uL22 and uL4 synthesis, respectively. Purple and blue triangles refer to repression of Rpf2 and Rrs1 synthesis, respectively.

The relative increase in the 40S peak (e.g., compare the profile in Figure 1 panels A and B) raised the possibility that this peak included uncommon precursor particles or degradation derivatives of 60S subunits. To address this, we determined the distribution of several r-proteins in sucrose gradients loaded with extracts from the P_gal_-uL4 cultures before and 8 hours after repression (Figure 1A-B). In both cases, the 40S peak contained the 40S protein uS4, but none of the 60S proteins tested (uL4, uL5 and uL18). We conclude that 60S-related particles did not co-fractionate with 40S subunits, indicating that the 40S peak contained only free 40S subunits.

For a more in-depth analysis of the 40S accumulation we ran sucrose gradients of lysates prepared from P_gal_-uL22, harvested at different times after the shift to glucose medium. If the accumulation of 40S subunits continues independently of the abolishment of 60S assembly, the ratio of 40S subunits to total ribosomal mass should increase throughout the experiment. Quantification of the gradient peaks shows that this ratio does in fact increase by 50-60% during the first couple of hours after repressing uL22 synthesis, but then becomes virtually constant (Figure 1K). Results from quantification of ribosomal peaks after repressing uL4, Rpf2, and Rrs1 fall on the same curve confirming that the results do not depend on how the 60S assembly is disrupted. Moreover, the fraction of ribosomal material in free 40S subunits increases sharply during the first couple of hours, then becomes constant, as expected because 40S subunits made in excess of 60S subunits cannot form 80S pairs (Figure 1L). The plateauing of the number of 40S relative to total ribosomes shows that either the 40S assembly stops after a few hours, or is offset by degradation of 40S subunits. As shown below, 40S assembly does indeed continue, so excess 40S must be degraded.

### Abrogating assembly of the 40S subunit does not inhibit 60S accumulation, but affects the structure of the 60S

We next tested the effect of preventing 40S subunit assembly by repressing the synthesis of early (eS1, uS4, uS7), and late 40S assembly proteins (uS10, uS11, eS31), as well as the 40S assembly factor Rrp7 (Figure 2 and Supplementary Figure 1C-I). The same pattern evolved regardless of how 40S assembly was abrogated: namely, the 40S peak was reduced and the 60S peak increased relative to the 80S peak. Although this increase was evident, the overlap between the 60S and 80S prevents an accurate quantification of these peaks (Figure 2A-C), but the conclusion is confirmed by quantification of ribosomal components (see below). Furthermore, the ratio between the ribosomal peaks and the A^260^ material at the top of the gradient decreased less after repression of 40S assembly than it did after repressing 60S assembly (compare Supplementary Figure 1 panels J-K with panels H-I). These results show that 60S assembly and accumulation continue during interruption of 40S assembly, a conclusion supported by our observation that GFP-tagged uL23 expressed from an inducible promoter 8 hours after repression of uS17 synthesis is incorporated into 60S subunits and 80S ribosomes (Supplementary Figure 2).

**Figure 2.**
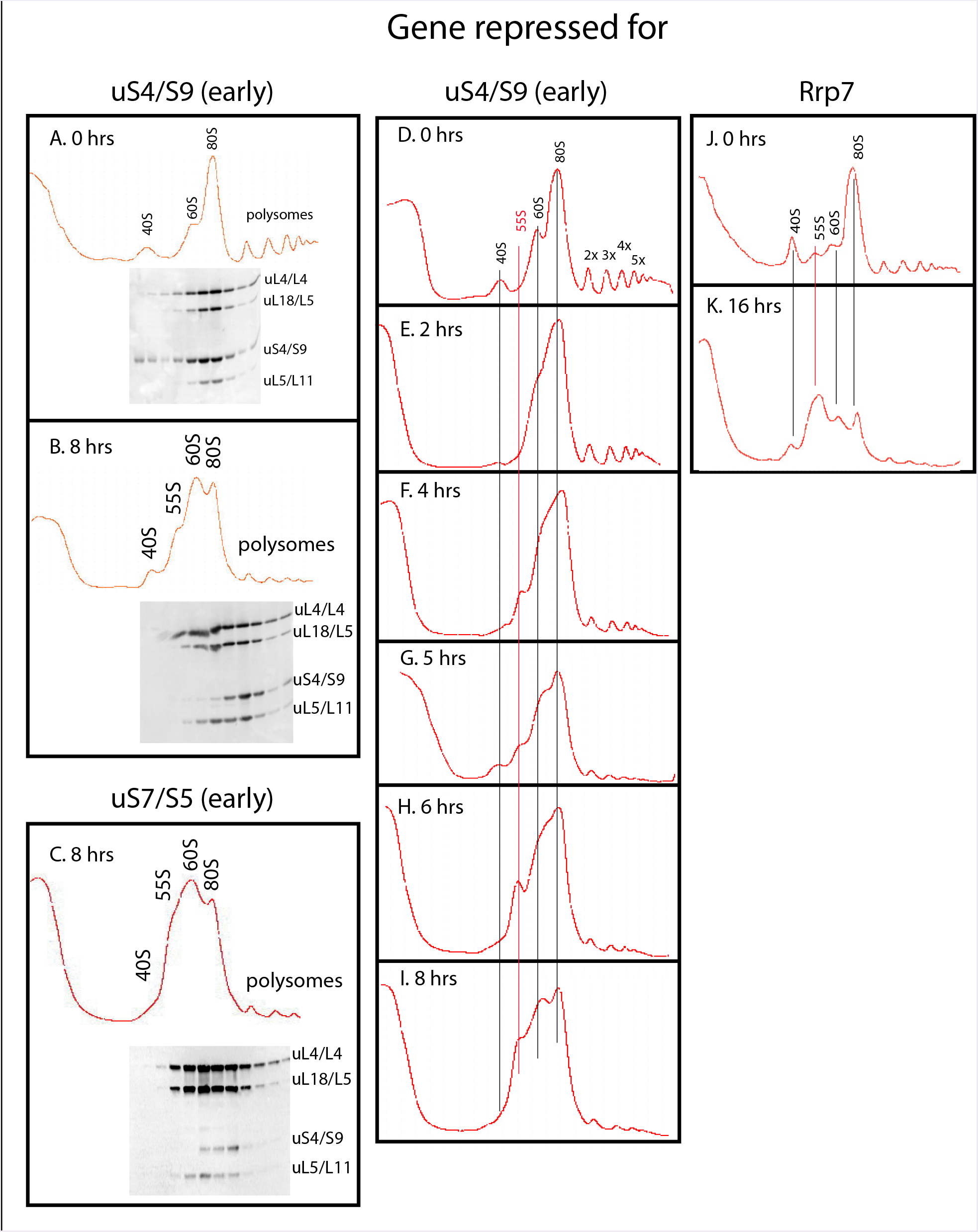
Sucrose gradient analysis of ribosomes after repressing the genes for the 40S r-proteins uS4 and uS7 and 40S assembly factor Rrp7. Cultures in YEP galactose medium were shifted to glucose medium for the indicated lengths of time before harvest. A. P_gal_-uS4 0 hours. B. P_gal_-uS4 8 hours. C. P_gal_-uS7 8 hours. D-I. P_gal_-uS4 0, 2, 4, 5, 6, and 8 hours, respectively. J-K. P_gal_-Rrp7 0 and 16 hours, respectively. For A-C, equal volumes of fractions in the ribosome portion of the gradient were analyzed for r-proteins uS4, uL4, uL5, and uL18 by western blot.

Although 60S accumulation continues in the absence of 40S production, we were intrigued to observe the build-up of a new peak, sedimenting slightly slower than 60S.

We refer to this new peak as “55S” (Figure 2, Supplementary Figure 3A-F). This particle has not previously been described, although it may be the reason for asymmetry of 60S peaks, trailing toward the top of the gradient, in several previous reports describing the effects of the depletion of 40S assembly factors (see e.g. (Milkereit et al. 2003; Perez-Fernandez et al. 2007)).

We followed the timing of the formation of this new peak by sucrose gradient analysis of extracts from cells harvested at different times after blocking uS4 synthesis (Figure 2D-I) or S31 synthesis (Supplementary Figure 3A-F). The 55S peak was not evident until 4-5 hours after the shift to glucose medium, but increased in size until 8 hours. No further increase was observed after 8 hours (compare Figure 2I with Supplementary Figures 1C and 3H and J). Thus, the 55S is apparently unstable such that an equilibrium between formation and degradation of 55S particles is established after about 8 hours. To determine if the accumulation of the 55S is affected by growth rates, we compared the height of the 55S peak after repressing the uS4 gene in cultures growing rapidly in rich YEP and relatively slower in synthetic medium (compare Figure 2D-I and Supplementary Figure 3 G-H with Supplementary Figure 3 panels I-J). The 55S peak was smaller compared to the 60S peak during the relatively slower growth in synthetic medium than in YEP medium, indicating that growth rate affects dynamics of 55S accumulation.

### The 55S particle is a derivative of the 60S subunit

Western analysis of the sucrose gradient fractions showed that the 55S peak contains large subunit proteins uL4, uL5, and uL18, but not the 40S protein uS4 (Figure 2A-C), suggesting that this novel peak is related to the 60S subunit. To test this conclusion we analyzed the RNA of the 55S peak. Extracts from Pgal-uS4 cells harvested before or 8 hours after repressing uS4 synthesis were centrifuged through sucrose gradients. RNA purified from ribosome-containing fractions of the 0-hour gradient was loaded in odd number slots of an agarose gel, and RNA from equivalent fractions of the 8-hour gradient was loaded in even number slots (Figure 3). After electrophoresis, RNA was transferred to a membrane and stained with methylene blue. The results show that the 55S peak contains 25S, but not 18S, rRNA (Figure 3A, lanes 8 and 10), confirming that the 55S peak is related to mature 60S. The 40S position contained 18S rRNA before (Figure 3A, lanes 3 and 5), but not after (lanes 4 and 6) repression of the uS4 gene, and the 80S peak contained both 18S and 25S rRNA both before and after (lanes 13-16).

**Figure 3.**
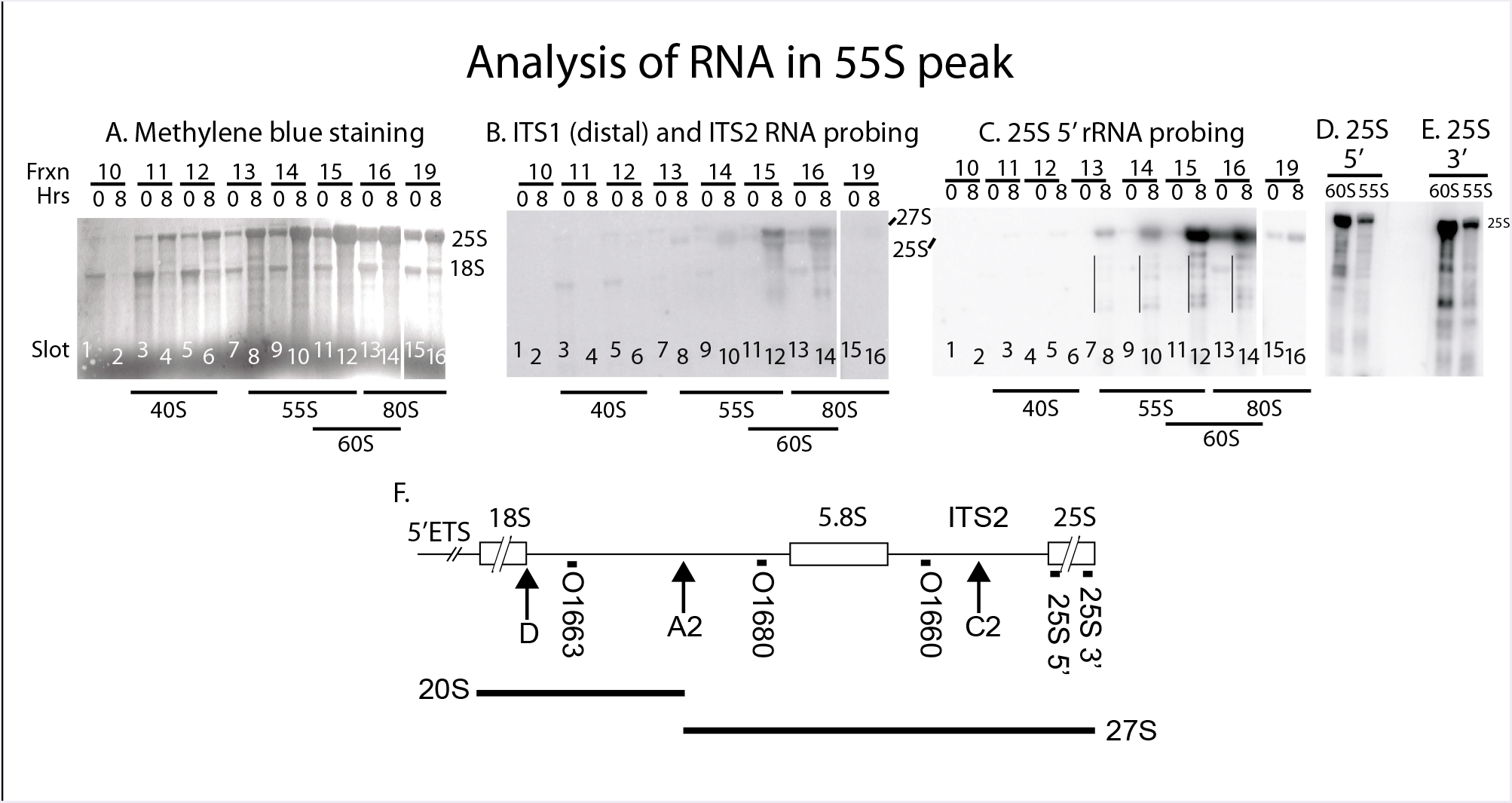
Analysis of RNA in the 55S peak appearing after repressing the gene for 40S r-protein uS4. P_gal_-uS4 was grown in YEP-galactose medium and aliquots were harvested at 0 hr and 8 hours after shift to glucose medium. Equal A^260^ units of whole cell extracts were fractionated on sucrose gradients. Samples of fractions from the 0 hour gradient were loaded in odd number slots of an agarose gel and samples of fractions from identical positions of the 8 hour gradient were loaded in even number slots. A-C. Analysis of rRNA in different ribosomal fractions. After electrophoresis, RNA was transferred to a membrane, which was stained with methylene blue (A), then probed with a mixture of equal radioactivity of the oligonucleotides hybridizing to ITS1 (distal to the A2 processing site) and ITS2 (B), stripped of radioactivity, and finally probed with an oligonucleotide specific to the 5’ end of 25S rRNA (C). The positions of the A^260^ peaks in the gradients are indicated below the images of the membrane. Degradation products of 25S rRNA are indicated by thin vertical lines. D-E. Northern blot of RNA from the 55S and 60S fractions of a sucrose gradient of samples harvested 8 hours after addition of glucose to a separate independent P_gal_-S4 culture and probed with radioactive oligonucleotides hybridizing to the 5’ (panel D) or the 3’ (panel E) ends of 25S rRNA. F. Map of the yeast Pol I rRNA transcription unit. The black boxes immediately below the map show the positions of the oligonucleotide probes used. The arrows labeled D, A2 and C2 indicate cleavage sites relevant for this study. Other processing sites are omitted. The major rRNA processing intermediates (20S and 27S) are indicated below the map.

To look for pre-25S rRNA processing intermediates we probed the blot with a mixture of oligonucleotides hybridizing to the distal part of ITS1 (O1680) and ITS2 (O1660) (Figure 3B and F), both parts of the 27S processing intermediates (Figure 3F). The 27S pre-rRNA band in the 8-hour gradient was much stronger for the 60S peak than for the 55S peak (Figure 3B, compare lanes 8 and 10 with lane 12). Pre-ribosomal intermediates accumulate in proportion to their average lifetime (Lindahl 1975), so the relatively weak 27S signal in the 55S peaks could be interpreted to mean that 55S is a short-lived precursor in the 60S subunit assembly pathway, while the 27S signal in the 60S peak would represent longer living precursors, probably the canonical 66S pre-60S particles (Woolford and Baserga 2013) that co-sediment with the mature 60S subunits under the conditions of our sucrose gradients. However, if this were the case, the amount of A^260^ material in the 55S peak should also be much smaller than the A^260^ material in the 60S peak, especially since most A^260^ material in the 60S peak represents mature 60S particles. In contrast to this prediction, the height of the 55S peak was at least half of the 60S peak (Figure 2B, C, H and I; Supplementary Figure 1C-M), arguing against the idea that the 55S is a 60S precursor particle. Therefore, the 55S must be a derivative of, rather than a precursor to, the 60S subunit. The reduction in the sedimentation coefficient presumably results mainly from a conformational change. Finally, the blot was stripped and probed with an oligonucleotide hybridizing to the 5’ end of mature 25S rRNA (Figure 3C and F). This showed that both the 60S and 55S peaks contained 25S rRNA fragments after, but not before, cessation of 40S assembly (marked with thin lines in Figure 3C), while 25S rRNA in the 80S was intact in both samples. The results in Figure 3 thus suggest that 60S subunits that fail to find a 40S partner become vulnerable to ribonuclease(s) and a conformational change.

To determine if the fragmentation of the 25S rRNA was due to exo- or endonucleases, we compared the patterns of bands generated by probing for the 5’ and 3’ ends of the mature 25S (Figure 3D-F). If the fragments in the 55S and 60S peaks were due only to digestion by a 3’>5’ exonuclease, such as the nuclear exosome, the 3’ probe should only hybridize to full length 25S rRNA. However, the 3’ probe hybridized to both full length and shorter fragments. Therefore, the 25S rRNA fragments cannot be formed exclusively by 3’>5’ exonuclease. Similarly, the fragments cannot be formed exclusively by a 5’>3’ exonuclease, since the 5’ probe hybridized to both full length and shorter 25S fragments. Thus, the fragmentation must involve endonucleolytic attacks.

Together the results from the uS4 repression experiments show that the 55S is a derivative of the 60S subunit that begins accumulating 4-5 hours after repressing the 40S assembly. After about 8 hours that ratio of formation and degradation of the 55S appears to be constant. Since the 55S is derived from the 60S subunit this suggests that the 60S subunit is destabilized, although its turnover is much slower than the degradation of the 40S subunits while 60S is blocked.

### The abundance of both 40S and 60S proteins decreases after inhibiting 60S protein synthesis, but abolishing 40S protein synthesis has little effect on 60S protein abundance

To further investigate the accumulation of ribosomal subunits, we measured the relative r-proteins concentrations (r-protein-x/total protein) after repressing an r-protein gene. Since disrupting ribosome biogenesis causes pleiotropic changes in the expression of numerous genes (Shamsuzzaman et al. 2017), we could not identify a protein suitable for western loading control. Instead, we compared intensities of r-proteins bands in the experimental samples to standard curves generated from a series of lanes on the same gel loaded with increasing amounts of a standard lysate (see Materials and Methods and Supplementary Figure 4 for details).

The relative concentration of ribosomal proteins does not change significantly in the parent strain (BY4741) after the shift to glucose (Figure 4A), as expected, since ribosome synthesis continues without interruption after the shift (Kief and Warner 1981). After repressing the 60S r-protein genes encoding uL4, uL11 or eL40, the relative concentration of 60S proteins uL4, uL5, and uL18 decreased (Figure 4B-D), as expected, since 60S assembly and accumulation are inhibited, while ribosomes made before the shift continue to make protein. Significantly, the relative concentration of uS4 <u>also</u> decreased, confirming that 40S accumulation is inhibited in response to cessation of 60S assembly.

**Figure 4.**
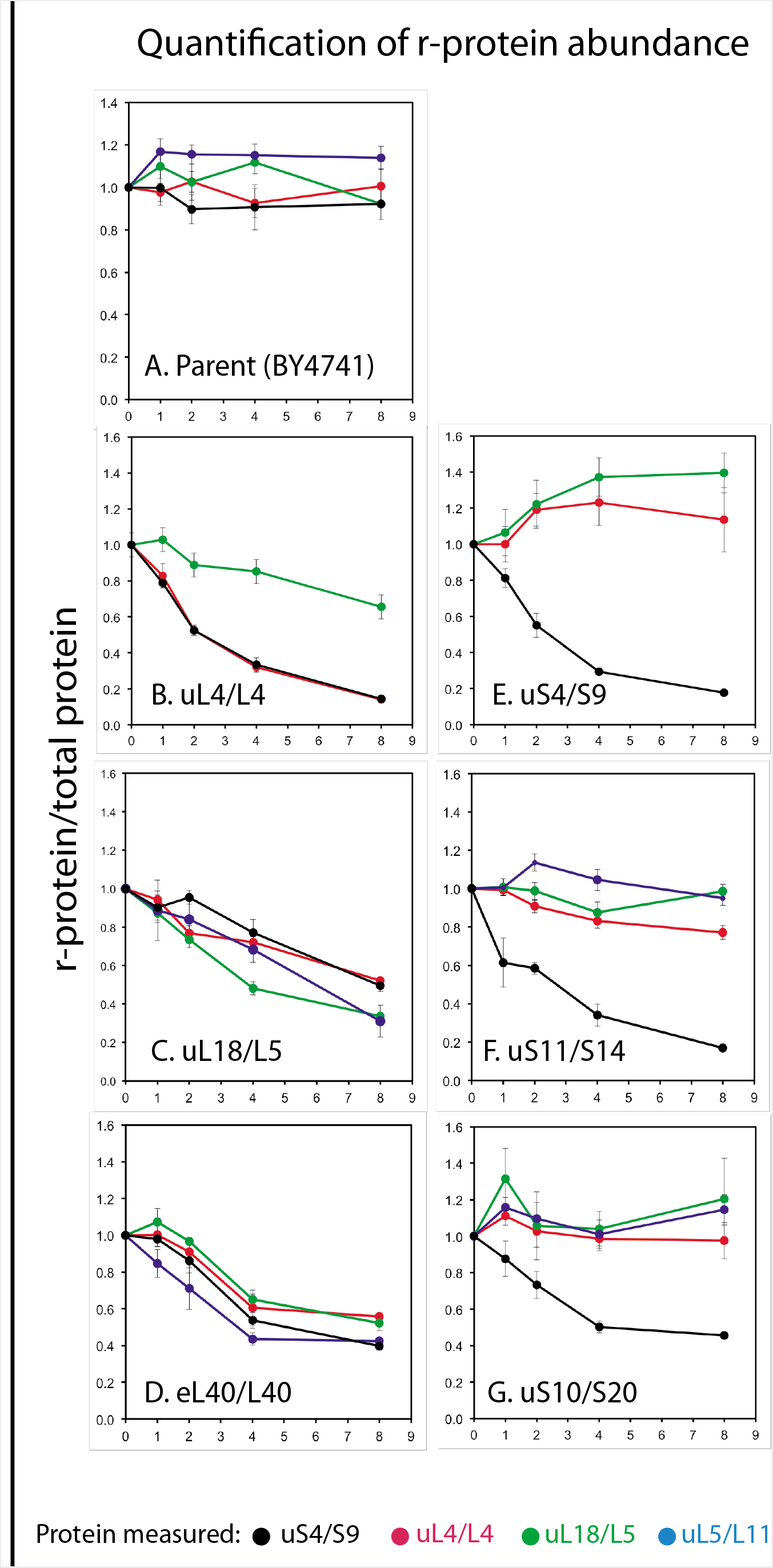
Quantification of r-proteins during repression of 40S and 60S r-protein genes. Strains in which the indicated r-protein genes are expressed from a galactose promoter, and the parent BY4741, were grown in YEP galactose. At time 0, the cultures were shifted to glucose medium and aliquots of the cultures were harvested at 0, 1, 2, 4, and 8 hours after the shift and used for western analysis of uS4, uL4, uL5, and uL18. The bands on the developed blots were quantified using a standard curve generated for each individual gel on the same blot as explained in the Materials and Methods and Supplementary Figure 4.

In contrast to the pan-subunit effect of repressing 60S protein synthesis, restraining the synthesis of the 40S proteins uS4, uS10, or uS11 generated a subunit-specific response, i.e. the specific concentration of uS4 protein declined, while the abundance of 60S proteins uL4, uL5, and uL18 changed little (Figure 4E-G). This result supports the conclusion that 60S accumulation is essentially unaffected when 40S assembly is perturbed. It is interesting that the uL18 concentration decreased slower in the uL4-repressed cells (Figure 4B) than in the eL40-repressed cells (Figure 4D). This may be related to the fact that uL18 is incorporated into an extra-ribosomal particle with 5S rRNA and other proteins, before it is transferred to the pre-60S ((Zhang et al. 2007), see Discussion).

In conclusion, the measurements of r-proteins relative to total protein confirm and extend the results from the sucrose gradient analysis: Inhibiting the synthesis of a 60S protein interferes with accumulation of <u>both</u> subunits, but abolishing the formation of 40S r-proteins only affects 40S accumulation.

### The 18S/25S rRNA ratio stays relatively constant after repressing 60S r-proteins genes, but declines after repressing 40S r-proteins genes

Finally, we analyzed rRNA transcripts. We first asked whether rRNA transcription continues after repression of r-protein genes. Since the transcribed spacers in the primary 18S-5.8S-25S rRNA transcripts are degraded both during normal subunit assembly and during turnover of ribosomal precursors, the abundance of transcribed spacer sequences should decline if no new transcripts are made. Accordingly, we measured the concentrations of ITS1 and ITS2 sequences in total rRNA transcripts.

RNA was purified from parent cells, Pgal-uS4 cells, and Pgal-uL4 cells before and at different times after the switch to glucose medium. Equal A^260^ units of total purified RNA from the cells were loaded onto a nylon membrane in a slot pattern, which was then probed with radioactive oligonucleotides complementary to the 20S part of ITS1 and ITS2 (Supplementary Figure 5, Figure 3F).

Figure 5A-B shows that the cell content of ITS1 and ITS2 sequences in the parent as well as in the Pgal-uS4 and Pgal-uL4 strains increased 2.5-3 fold after the addition of glucose. This increase may be due to a decreased rate of degradation of ITS1 and ITS2. In any case, the experiment shows that the abundance of ITS sequences does not decline, indicating that rRNA transcription continues after repression of r-protein synthesis. This conclusion is supported by run-on transcription measurements of RNA polymerase occupancy on rRNA genes. Cell extracts were prepared before and at 6 and 15 hours after repressing uL4 synthesis under conditions that preserve transcribing RNA polymerase molecules on their template. The extracts were then incubated in the presence of radioactive UTP to allow RNA polymerase I to complete their current round of transcription, but not to initiate new rounds of transcription. The radioactive run-on products were hybridized to a membrane loaded with denatured pDK16 plasmid DNA containing the full Polymerase I transcription unit (Lindahl et al. 1994). The results show that the intensities of the rRNA run-on transcripts are either unchanged or increased after the shift (Figure 5C). Taken together, the results presented in Figure 5A-C demonstrate that rRNA transcription continues, even though the synthesis of an essential r-protein is repressed.

**Figure 5.**
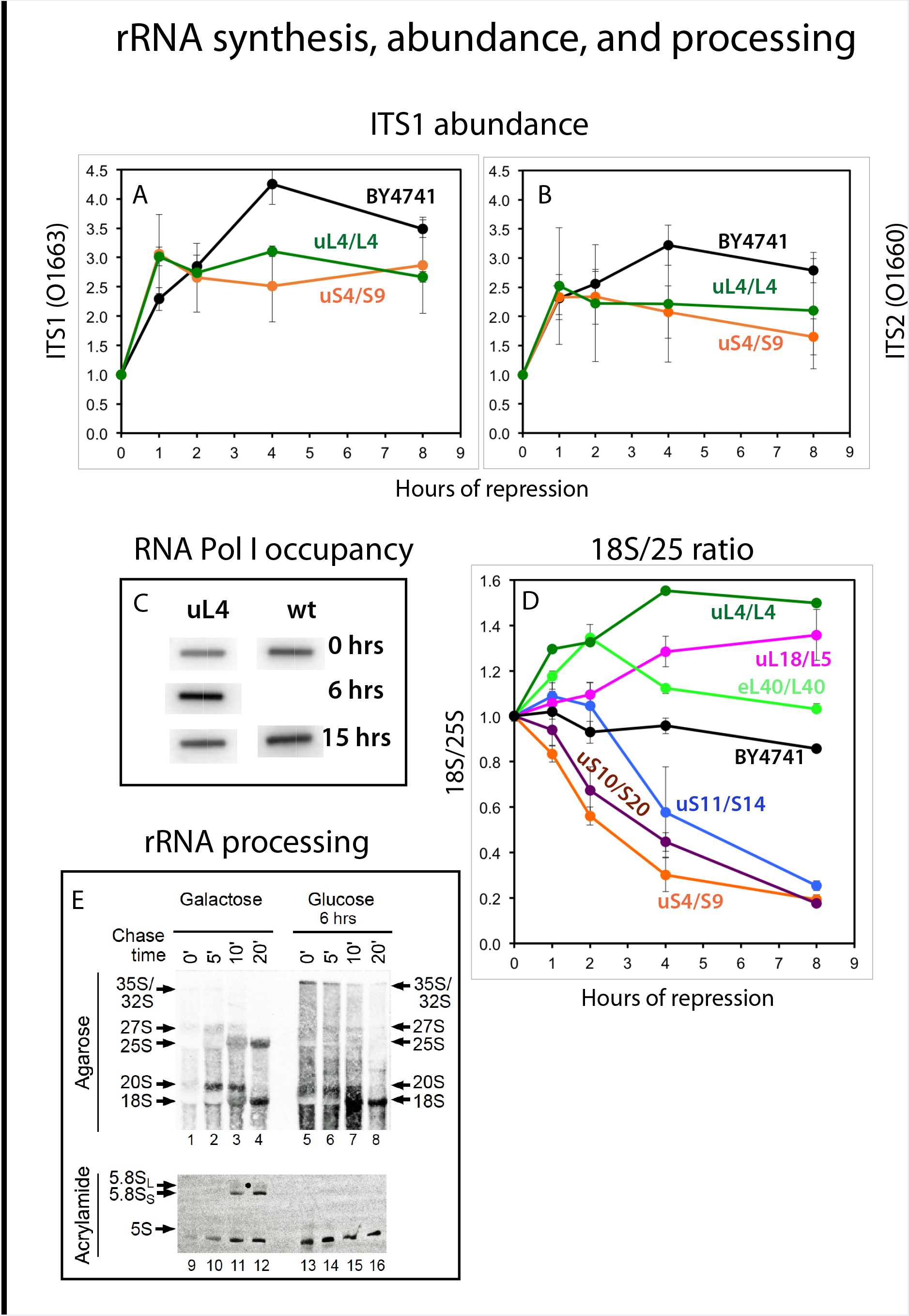
Analysis of rRNA after repression of 40S and 60S r-protein genes. Strains in which the indicated r-protein genes are expressed from a galactose promoter, and the parent BY4741, were grown in YEP galactose (panels A, B, and D) or synthetic complete galactose medium (panels C and E). At time 0, the cultures were shifted to glucose medium. Aliquots of the culture were harvested at 0, 1, 2, 4, and 8 hours after the shift. A-B. Measurement of transcripts containing ITS1 and ITS2. Total RNA was purified and equal A^260^ units were loaded onto a nylon membrane in a slot format. The membrane was probed with ITS1 oligonucleotide O1663 (A) or ITS2 oligonucleotide O1660 (B); see Figure 3F for the positions of the rRNA transcription to which the probes hybridize. The washed membranes were scanned and bands were quantified. C. Run-on analysis of rRNA transcription during repression of uL4 synthesis. Aliquots of cultures of Pgal-uL4 in synthetic medium were harvested before and 6 hours after shift to glucose medium, lysed and used for run-on labeling with α-^32^ATP. The products were hybridized to slot blots of pDK16 carrying the 35S rRNA transcription unit, but not the “non-transcribed spacer” and the 5S rRNA gene (Lindahl et al. 1994). D. Ratio between accumulated 18S and 25S rRNA. Total RNA was prepared from the indicated Pgal-x cultures and fractionated on agarose gels. After electrophoresis, the gels were stained with ethidium bromide and photographed. The images were then scanned and the intensities of the 18S and 25S bands were quantified. E. Pulse-chase analysis of rRNA processing during repression of uL4 synthesis. Pgal-uL4 was grown in synthetic medium and harvested 0 or 6 hours after shift to glucose medium. Cells were labeled with ^3^H-uracil for three minutes and chased with a large excess of non-radioactive uracil for the indicated times. Total RNA was extracted and analyzed by agarose gel electrophoresis.

Having established that rRNA synthesis continues after repression of r-protein genes, we analyzed the relative abundance of 18S and 25S rRNA by fractionating total purified RNA on agarose gels and quantifying ethidium bromide-stained bands. As expected, the ratio between 18S and 25S rRNA did not change after shifting the parent strain (BY4741) to glucose medium (Figure 5D). After repression of 40S protein genes, the ratio declined and was ultimately decreased 4- to 5-fold, in accordance with the finding that the abundance of 40S subunits and 40S r-proteins declines, while the abundance of 60S subunits and proteins remains relatively constant. In contrast, the 18S/25S ratio increased 20-50% after repression of 60S r-protein genes, then either stabilized or declined back to 1:1. The relatively minor changes in the ratio of 18S/25S rRNA after repression of 60S r-protein synthesis in comparison with the change during inhibition of 40S assembly also supports the conclusion that accumulation of both subunits is inhibited by repression of 60S r-protein genes. The subtle variances in the kinetics of the 18S/25S ratio after the abolition of different proteins from the same subunit are most likely due to differences in the rate of turnover of abortive assembly intermediates depending on which step of the assembly is affected by the cessation of individual r-proteins.

Since 40S accumulation is inhibited after disruption of 60S assembly, we asked if this is due to inhibition of 40S assembly and 18S rRNA maturation or to degradation of 40S subunits after assembly. To answer this we followed rRNA processing before and after cessation of uL4 synthesis. Total RNA was pulse labeled with ^3^H-uracil for 2 minutes at which time a large excess of non-radioactive uracil was added. Total RNA prepared before (0) and 5, 10, and 20 minutes after addition of an excess of non-radioactive uracil was fractionated by agarose gel electrophoresis and bands were visualized by fluorography (Figure 5E). Prior to repression of uL4 synthesis we observed the expected pattern, i.e. radioactive rRNA was first seen in the 35S transcript then moved to 27S and 20S processing intermediates and finally to mature 25S, 18S rRNA and 5.8S rRNA over a 10-20 minute period (lanes 1-4 and 9-12). Six hours after repression of uL4 synthesis no mature 25S rRNA or 5.8S rRNA was formed (lanes 4-8 and lanes13-16) in accordance with the expectation that 60S ribosomal components are degraded when 60S assembly is prevented. However, 18S was still matured (compare lanes 5-8 with lanes 1-4). These results are consistent with the finding that 18S processing continues, but 25S processing does not, after temperature inactivation of the 60S assembly factor Rrp1 (Horsey et al. 2004). The subunit-specific effect on the processing of rRNA also concurs with the fate of r-proteins during inhibition of rRNA assembly: newly synthesized 60S proteins, but not 40S proteins, rapidly turn over after inactivation of a 60S assembly factor (Gorenstein and Warner 1977). The dichotomy between the processing of 18S rRNA and the blockage of 40S subunit accumulation after cessation of 60S assembly leads to the conclusion that the interference with 40S accumulation occurs after assembly of a 40S particle containing 18S rRNA processed to its final length.

Interestingly, 5S rRNA processing is not affected by the disruption of 60S assembly (compare lanes 9-12 with lanes 13-16). We have previously observed that inactivation of the endonuclease RNase MRP stops processing of both the 18S and 25S/5.8S segments of the 35S precursor rRNA thereby blocking maturation of 18S, 5.8S and 25S rRNA, but also does not affect 5S rRNA accumulation (Lindahl et al. 2009). Possibly, 5S rRNA maturation is independent of 60S assembly because the 5S rRNA is initially incorporated into an extra-ribosomal particle with r-proteins uL5 and uL18 and assembly factors Rpf2 and Rrs1 (Zhang et al. 2007).

### Constraining 60S assembly inhibits nuclear export of both pre-40S and pre-60S particles, but 40S assembly inhibition affects only pre-40S export

Ribosomal precursor particles are exported from the nucleus to the cytoplasm once they become competent to bind nuclear export factors (Fischer et al. 2015; Malyutin et al. 2017). If assembly is blocked r-proteins accumulate in the nucleus (Hurt et al. 1999; Milkereit et al. 2003). To determine if the disruption of 40S or 60S assembly affects nuclear export of subunit precursors, we introduced low copy plasmids carrying uS5-GFP or uL23-GFP fusion genes expressed from the native promoters into the Pgal-eS31 and P_gal_-eL43 strains, and performed confocal microscopy of the resulting strains 16 hours after shift to glucose medium. Figure 6 shows that repression of the eS31 gene results in nuclear accumulation of uS5-GFP, but not of uL23-GFP, confirming that preventing assembly of the pre-40S precursor to an export-competent stage specifically traps 40S protein in the nucleus. As also expected, uL23 also accumulated in the nucleus after blocking pre-60S assembly by repressing the eL43 gene. It was, however, surprising, that repression of the eL43 gene <u>also</u> caused nuclear accumulation of uS5. Thus, abolishing 60S assembly inhibits export of <u>both</u> subunits.

**Figure 6.**
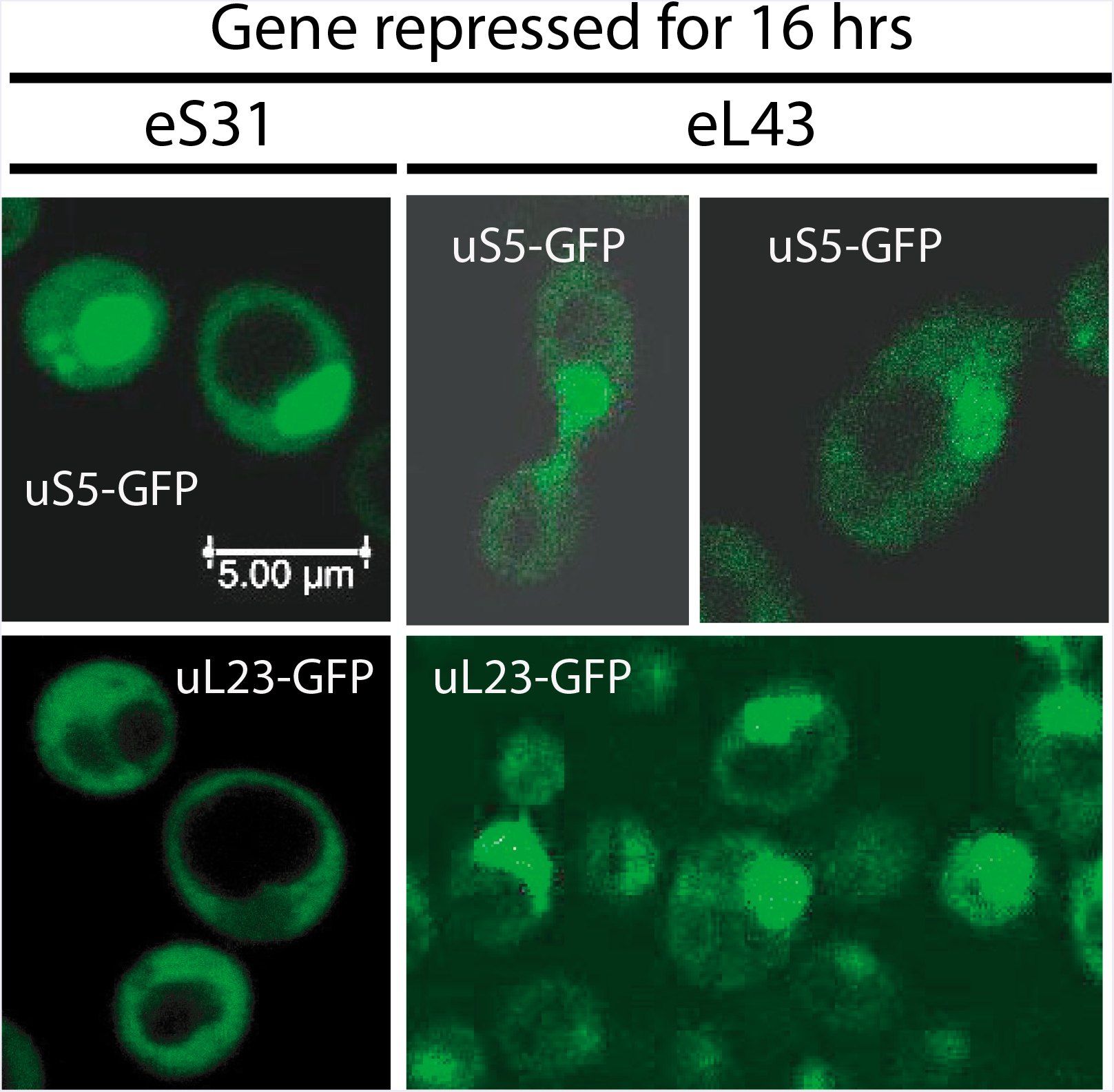
Cellular localization of r-proteins during repression of a 40S or a 60S r-protein gene. Pgal-eS31 or Pgal-eL43, each carrying plasmid-borne genes for uS5-GFP or uL23-GFP expressed for the native promoters, were grown in synthetic galactose medium and shifted to synthetic glucose medium for 16 hours. Cells were inspected by confocal microscopy. A uniform distribution of the GFP-tagged protein between the nucleus and cytoplasm indicates that the export of ribosomal precursor particles carrying the tagged protein is normal. A buildup of the GFP-tagged in the nucleus indicates that ribosomal precursor particles carrying the tagged protein are not matured to nuclear export competency.

### Vacuole expansion

In agreement with previous observations (Bernstein et al. 2007), Figure 6 also shows that extremely large vacuoles develop when assembly of either subunit is constrained. We speculate that this may be the result of massive turnover of ribosomal components when complete assembly of ribosomal subunits is inhibited.

## Discussion

### Disruption of assembly of one subunit affects the stability of the other

The large and small ribosomal subunits are formed through independent pathways that originate from splitting a common precursor rRNA transcript. During undisturbed ribosomal assembly this assures production of equal numbers of the subunits. However, mutations in genes for ribosomal components or one of the numerous assembly factors can inhibit assembly of one subunit and thus distort the balance between the numbers of subunits. Here we have investigated whether there are mechanisms that can counteract departures from the 1:1 ratio of the subunits. Our extensive quantification of ribosomal subunits as well as their rRNA and protein components after repressing the assembly of one of the subunits shows that disrupting the formation of the 60S subunit impedes the accumulation not only of the 60S, but also of the 40S subunit (Figures 1K-L, 4B-D, and 5D). The converse is not the case: abolishing the assembly of the 40S subunit does not prevent accumulation of 60S subunits (Figures 2, 4E-G, and 5D), but it causes fragmentation of the 25S rRNA in these subunits and formation of a 55S particle that is derived from the 60S subunit (Figure 3).

### Post-assembly turnover of the 40S subunit

When a gene encoding an essential r-protein is repressed, assembly of the subunit to which the protein belongs cannot be completed. However, transcription of rRNA (Figure 5 A-C), and r-protein genes continues (Horsey et al. 2004; Thapa et al. 2013; Shamsuzzaman et al. 2017). Since assembly of the two ribosomal subunits occurs along separate largely independent pathways (Woolford and Baserga 2013), we were surprised that inhibition of the 60S subunit prevents accumulation of the 40S subunit. To understand why 40S accumulation is linked to 60S assembly, we tested if 40S particles are assembled in the absence of formation of 60S subunits. Pulse-chase experiments showed that the 18S rRNA is processed to its final size (Figure 5E), suggesting 40S subunit precursors are indeed assembled. Therefore the cessation of 40S accumulation must be due to degradation after assembly. However, excess 40S relative to total ribosomal material initially accumulates showing that it takes a few hours after cessation 60S assembly before 40S turnover becomes noticeable. The delay in the turnover could have different causes: A pool of free 40S subunits may have to build up before the chance of encountering a degradation enzyme becomes significant, or synthesis of degradation enzyme(s) must be induced.

The final cleavage of pre-18S rRNA (20S) that brings 18S to its final length separates the 18S moiety from the proximal part of ITS1. This cleavage occurs in the cytoplasm in cytoplasmic pre-40S-60S complexes, i.e. after export to the cytoplasm (Lebaron et al. 2012; Strunk et al. 2012; Garcia-Gomez et al. 2014). Nevertheless, the distribution of r-proteins shows that export of pre-40S is inhibited during repression of 60S protein synthesis (Figure 6). It was shown previously that the concentration of ribosomal proteins is similar in the nucleus and cytoplasm during normal growth, but inhibition of the maturation of one of the subunits cause concentration in the nucleus to exceed the concentration in the cytoplasm (Hurt et al. 1999; Milkereit et al. 2003). In agreement with this, we also observed nuclear accumulation of uS5 after repression of the eS31 gene (Figure 6). Unexpectedly, both the 60S protein uL23 and the 40S protein uS5 accumulated in the nucleus after repression of the uL40 (Figure 6) or uL4 genes (not shown), showing pre-40S export is linked, at least to some degree, to 60S assembly. Perhaps pre-60S particles are necessary for complete modification rRNA and/or proteins in the 40S subunit, and under-modification could affect the stability and/or export nuclear pre-40S particles. It is also known that mature 40S subunits containing mutated rRNA are degraded (Rodriguez-Galan et al. 2015), but there are no known mutations in the 18S sequence of our strains. We speculate that 40S subunits accumulating as free (non-60S bound) particles due to the deficit of 60S subunits could be vulnerable to cytoplasmic nucleases, because the interface region, which is rich in rRNA loops, is exposed in the free subunits, but protected while it is paired with a 60S subunit.

In summary, our experiments suggest that during abrogation of 60S assembly, some of the pre-40S may be turned over in the nucleus, but most of the 40S degradation likely occurs in the cytoplasm. The coupling of 40S accumulation to 60S maturation could thus be the result of successive processes in both the nucleus and the cytoplasm.

### 60S subunits are destabilized when 40S assembly is blocked

Repression of 40S r-protein genes leads, with a delay of several hours, to fragmentation of a significant fraction of the 25S rRNA in free 60S subunits and to the appearance of a novel ribosomal particle (55S, Figure 2) derived from the 60S subunit. Thus, the 60S subunit is also destabilized during abrogation of 40S assembly, but the turnover rate of 60S is much lower than the degradation of 40S. This attack on the 60S most likely happens in the cytoplasm, because 60S proteins do not accumulate in the nucleus after inhibition of 40S assembly (Figure 6), suggesting that 60S assembly and nuclear export proceeds unimpeded. Some of the fragmented rRNA in the 60S peak hybridizes to the transcribed spacer probes. Processing of 25S rRNA is believed to be complete prior to nuclear export, but precursor 60S particles containing ITS2 can escape to the cytoplasm during impaired ITS2 processing (Sarkar et al. 2017). We speculate that nuclear processing of pre-25S rRNA may be subtly distorted in the absence of 40S assembly, even though the 40S and 60S assembly pathways are believed to be largely independent. Cytoplasmic pre-60S particles form defective 80S ribosomes that are degraded (Sarkar et al. 2017), but it is conceivable that they can also be degraded if they fail to form 80S couples due to the deficiency in the number of 40S subunits. As suggested above for the 40S subunit, the 60S interface side could be vulnerable to nucleases in the absence of the 40S partner. This interpretation is supported by the fact that the 25S rRNA in 80S appears to be intact (Figure 3C). Another possibility is that the 60S may be under-modified in the absence of 40S assembly and therefore subject to degradation, e.g. by the pathways that eliminate 60S particles with mutations in the peptidyl transferase center (Cole et al. 2009).

### Variation in the rates of degradation of uL18 after repression of different 60S r-protein genes

The uL18 concentration decreases more slowly during repression of uL4 synthesis than during inhibited eL40 gene expression (Figure 4B and D). This may be related to uL18 forming a complex with uL5, 5S rRNA, and two 60S assembly factors prior to its incorporation into pre-60S particles late in the assembly process (Zhang et al. 2007). Repressing the uL4 gene aborts the 60S assembly at an early step of the assembly pathway (Gamalinda et al. 2014), which may prevent transfer of uL18 to a pre-60S particle. Therefore, uL18 may remain in the pre-incorporation complex where it may be semi-protected from rapid degradation. In contrast, eL40, like uL18, is a late assembly protein, so interrupting the eL40 synthesis may stop assembly <u>after</u> uL18 has been transferred to the pre-60S particle and thus make uL18 as vulnerable as the rest of the pre-60S particle.

### Implications

We hypothesize that the linking of 40S accumulation to 60S subunit formation has evolved to secure proper function of the translation process. A surplus of 40S subunits would lead to formation of translation initiation complexes that cannot be converted to translating 80S ribosomes because of the shortage of 60S subunits. Potentially, this could indiscriminately sequester mRNAs and distort the synthesis of many proteins. Excess free 60S subunits would not have a similar effect on translation.

Our findings may also be relevant to diseases caused by mutations in genes for r-protein and factors for ribosome assembly and rRNA modification (Mattijssen et al. 2010; Narla and Ebert 2010; Tafforeau et al. 2013; Danilova and Gazda 2015; Farley and Baserga 2016; Bustelo and Dosil 2018). Since ribosomal assembly in yeast shares many aspects of human ribosome formation (Tafforeau et al. 2013), our work suggests that the mechanism for such diseases may involve interactions between subunit assembly and stability. Experimental protocols involving many hours of inhibition of ribosomal assembly are occasionally criticized, because this allows development of secondary reaction not directly related to the initial assault (e.g. (Kos-Braun and Kos 2017)). However, in the context of genetic diseases, the mutational stress is permanent, making it relevant to investigate the long time effect of disturbing ribosome assembly.

Ribosome formation requires a substantial fraction of the cell resources. During normal growth the fraction of the cell mass constituted by ribosomes increases with growth rate, which has led to the dogma that ribosome formation is optimized for minimal drain on cell resources (Maaløe and Kjeldgaard 1966; Warner 1999). Indeed, signaling pathways emanating from TOR (Target Of Rapamycin) repress transcription of ribosomal genes during poor nutritional conditions or oxidative stress (Claypool et al. 2004; Mayer and Grummt 2006; Philippi et al. 2010). However, our results show that preservation of resources is not always the cell’s priority. First, transcription of rRNA and r-protein genes continues unabated when assembly of one of ribosomal subunits is abrogated. Second, excess 40S subunits are turned over after resources have been expended on their assembly and maturation. Similarly, ribosomal components are synthesized, but turned over, when processing enzymes required for assembly of one or both subunits are inactivated (Gorenstein and Warner 1977; Horsey et al. 2004; Lindahl et al. 2009). In all cases there is a massive turnover of ribosomal material rather than a reduction in synthesis and assembly of ribosomal components, suggesting that TOR cues to reduce transcription of ribosomal genes are not activated in these situations.

This notion is supported by the fact that repression of uL4 synthesis did result in accumulation of a 23S rRNA transcript (Figure 5E) similar to the one observed when nutritional conditions or oxidative chemicals deactivate TOR (Kos-Braun and Kos 2017). We suggest that TOR is only deactivated by stress originating from sources external to the cell, but not by stress emanating from internal sources such as mutations in gene for ribosomal components or assembly factors.

## Materials and Methods

### Strains and growth conditions

Yeast strains were described previously (Ferreira-Cerca et al. 2005; Pöll et al. 2009; Thapa et al. 2013). In each strain a gene encoding a r-protein or ribosome assembly factor was expressed from the *Gal1/10* promoter. Many r-proteins in yeast are encoded by two different alleles that in most cases differ by a few amino acids. For our studies, in cases where two alleles exist in wildtype yeast, the second allele was deleted such that synthesis of the gal-controlled protein is repressed in glucose medium. With the exception of r-protein uL4, we only analyzed expression of one of the two alleles from *GAL1/10;* see (Thapa et al. 2013) for identification of the allele used. However, since abrogating the synthesis of different proteins from a given subunit generated the same pattern of ribosomal subunit accumulation, it is unlikely that the particular r-protein allele used has any important effect on the results. For repression of uL4 synthesis, all experiments, except in Figure 5 C and E, were done with JWY8402 expressing the gene encoding uL4A (RPL4A) from the galactose promoter. In Figure 5C and E, the gene encoding uL4B allele (RPL4B) was under galactose control (strain YLL2083).

Cultures were grown in YEP-galactose (1% yeast extract, 2% peptone, 2% galactose) or galactose synthetic complete medium lacking uracil with shaking at 30° (Sherman et al. 1979). Once cell density reached an OD^600^ of 0.5-0.8 corresponding to about 0.6-1x10^7^ cells per ml, the culture was shifted to pre-warmed YPD medium (1% yeast extract, 2% Bacto Peptone and 2% glucose) or glucose synthetic complete medium lacking uracil. Cultures were diluted with pre-warmed medium whenever necessary to keep the OD^600^ below 0.8. Cell density was measured in a 10 mm cuvette using a Hitachi U1100 spectrophotometer (Hitachi High-Technologies Corporation, Japan).

Plasmids carrying fusions of genes for GFP and uS5 or uL23 expressed from the native r-protein promoters were also described previously (Hurt et al. 1999; Milkereit et al. 2003). To enable induction of uL23-GFP synthesis during repression of uS17 synthesis, a gene for the artificial beta-estradiol-sensitive transcription factor Z3EV was introduced into Pgal-uS17, and the uL23-GFP gene was placed under control of the Z3EV-reponsive promoter (McIsaac et al. 2013).

### Western analysis

Cells were spun down at 5K for 10 minutes before being washed with and suspended in Buffer A (20 mM Tris-HCl, 150 mM KCl, 5 mM Na-EDTA, 0.1% Triton X-100, 10% glycerol), dithiothreitol (1 mM) and phenylmethylsulfonyl fluoride (20 μl/ml). Cells were lysed by vortexing for 15 minutes at 4°C with glass beads. A^280^ measurements were used to calibrate the amount of lysate applied to 15% polyacrylamide gels. After electrophoresis, proteins were transferred onto PVDF membranes for 1.5 hours at 130V using the Trans-blot Cell apparatus (Bio-Rad). The membranes were then incubated in a blocking solution (20 mM Tris, pH 7.5 and 150 mM NaCl, 0.1% Tween 20, dry 5% milk (LabScientific, Inc.)) before incubating with primary antisera (1:10000-1:4000 dilutions). Rabbit polyclonal antisera for the yeast ribosomal proteins were prepared for our laboratory by Covance (Princeton New Jersey, USA) using synthetic peptides with the sequence of 20-22 N-terminal amino acids of uS4, uL4, uL5, and uL18 as antigens. The membrane was cleared of excess antisera washing buffer (25 mM Tris-Cl (pH 7.4), 2.5 mM KCl, 200 mM NaCl) before it was incubated with Goat Anti-Rabbit IgG (H+L)-AP Conjugate secondary antibody (Bio-Rad). Bands were detected by exposing the membrane to Amersham ECF substrate (GE Healthcare) followed by scanning on Storm 860 Imager System (Molecular Dynamics). A standard curve was prepared on each blot by loading increasing amounts of a whole cell lysate from the parent strain (BY4741) into consecutive gel slots of the SDS-polyacrylamide gel. The remaining slots in each gel were loaded with a constant number of A^280^ units from a Pgal-protein-x strain harvested before and after addition of glucose (Supplementary Figure 4). After probing the western blot with a cocktail of the indicated antisera, standard curves were constructed for each antiserum using the band intensities of the BY4741 lanes. The correlation coefficients for the standard curves were >0.95, corroborating the accuracy of the method (Supplementary Figure 4).

### Sucrose gradient analysis

Many reports on ribosome sucrose gradients describe adding cycloheximide to the growing culture before harvest. However, cycloheximide artificially increases the polysome content even after short incubations with the drug (Helser et al. 1981; Panasenko and Collart 2012) (and our unpublished experiments). Accordingly, we did not add cycloheximide to the culture, but did include it in the lysis buffer to preserve polysomes after lysis. Cells were quick-chilled over ice, spun down, and washed with ice-cold water. The pellet was then resuspended in ice-cold gradient buffer (50 mM Tris Acetate pH7, 50 mM NH_4_Cl, 12 mM MgCl_2_, 1 mM DTT) containing 50 μg cycloheximide per ml. The mixture was transferred to a tube (Sarstedt NC9437081) containing 2.5 g glass beads (Sigma G9268 or G8772) and vortexed five times at maximum speed for thirty second intervals at 4°C. The resulting cell lysate was spun down twice at 13,000 g for 15 minutes at 4°C, and the pellet was discarded after each spin. Equal A^260^ units of the supernatant after the second spin were loaded onto 10-50% sucrose gradients in gradient buffer. The gradients were spun at 40,000 rpm for 4 hours at 4°C using a SW40Ti Beckman rotor. Fractions (500 μl) were collected using the ISCO Foxy Jr fraction collector, pumping at 1 ml/min. The relative amounts of A^260^ material in the different peaks were determined using ImageJ.

## RNA analysis

*Total RNA extraction*. Cells were harvested by centrifugation and stored at -80°C. The cells were then resuspended in ice-cold water and RNA was extracted using the phenol chloroform method as described (Lindahl et al. 1992).

*Extraction of RNA from RNA polysome gradient fractions* was performed as described (https://case.edu/med/coller/Coller%20Protocol%20Book.pdf) (Cigan et al. 1991; Nielsen et al. 2004). Briefly, individual sucrose gradient fractions were mixed with an equal volume of ethanol and precipitated overnight at -80°C. The pellets were resuspended in LET (25 mM Tris-HCl pH 8.0, 100 mM LiCl, 20 mM Na-EDTA) and 1% sodium dodecylsulfate, extracted twice with one volume phenol/chloroform/isoamyl alcohol (25:24:1), and precipitated overnight at 80°C using 2.4 volumes ice-cold ethanol and 1/5 volume 10 M ammonium acetate.

*Ribosomal RNA analysis*. (i) The ratio between 18S and 25S rRNA was measured by fractionating 3 μg of total RNA on 0.8 or 1.2% agarose gels that was stained with ethidium bromide after the run. (ii) ITS1 and ITS2 were quantified by “slot blot analysis” as follows. Three µg total RNA in sterile water was deposited directly onto an Amersham Hybond-N nylon membrane (GE Healthcare Life Sciences) using the Minifold II slot blot system (Schleicher & Schuell). After cross-linking, membranes were hybridized with ^32^P-end-labeled probes (Sambrook et al. 1989) in a solution made from 20 mL 100X Denhardt’s solution (http://cshprotocols.cshlp.org/content/2008/12/pdb.rec11538.full?textonly=true), 60 mL 20x SSC, 120 mL H_2_O, and 2 mL 10% SDS. The membrane was incubated with rotation in the solution for at least 1 hour. Radioactive probes were added (5x10^6^ cpm per chamber) after which the mix was incubated with rotation at 37° overnight. Finally, the blot was washed three times for 10 minutes at room temperature with 6xSSC and 0.01% SDS and exposed to a storage phosphor screen (Molecular Dynamics) for 4 hours. Bands were detected by scanning on a Storm 860 Imager System or a Typhoon 9200 Imager (Molecular Dynamics). (iii) RNA purified from sucrose gradient fractions was fractionated by gel electrophoresis on a 0.8% agarose gels in 0.5xTBE (45 mM Tris-borate, pH8.3, 1 mM EDTA-Na) and was transferred onto an Amersham Hybond-N nylon membrane (GE Healthcare Life Sciences) using a Model 230600 Boekel Vacuum Blotter for 2 hours at ~ 40 mbar. The RNA was then crosslinked to the membrane and stained with 0.04% methylene blue in methanol. Next, the membrane was destained with 25% ethanol and hybridized with ^32^P-end-labeled probes as above.

Sequences of the oligonucleotide probes used are: O1663

CTCTTGTCTTCTTGCCCAGTAAAAG; O1660 AGGCCAGCAATTTCAAGTTAACTCC. O1680: CCAGTTACGAAAATTCTTGTTTTTGAC; 5’ 25S:

ACTCCTACCTGATTTGAGGTCAAACC; 3’ 25S: GGCTTAATCTCAGCAGATCGTAAC. See also Figure 3F.

### Other methods

The bands on northern and western images were quantified and analyzed using ImageJ software. Run-on transcription was performed as described (Elion and Warner 1986). Pulse-chase labeling was performed essentially as described by (Dunbar et al. 2000). Error bars indicate the Standard Error of the mean based on samples from three independent cultures (biological repeats) and two measurements of each sample (technical repeats).

## Acknowledgments

This work was supported by grant 0920578 from the National Science Foundation and a grant from UMBC. We thank John Woolford (Carnegie Mellon University), Philipp Milkereit, (University of Regensburg), and Ed Hurt (The University of Heidelberg) for strains.

---

**Supplementary Figure 1.**
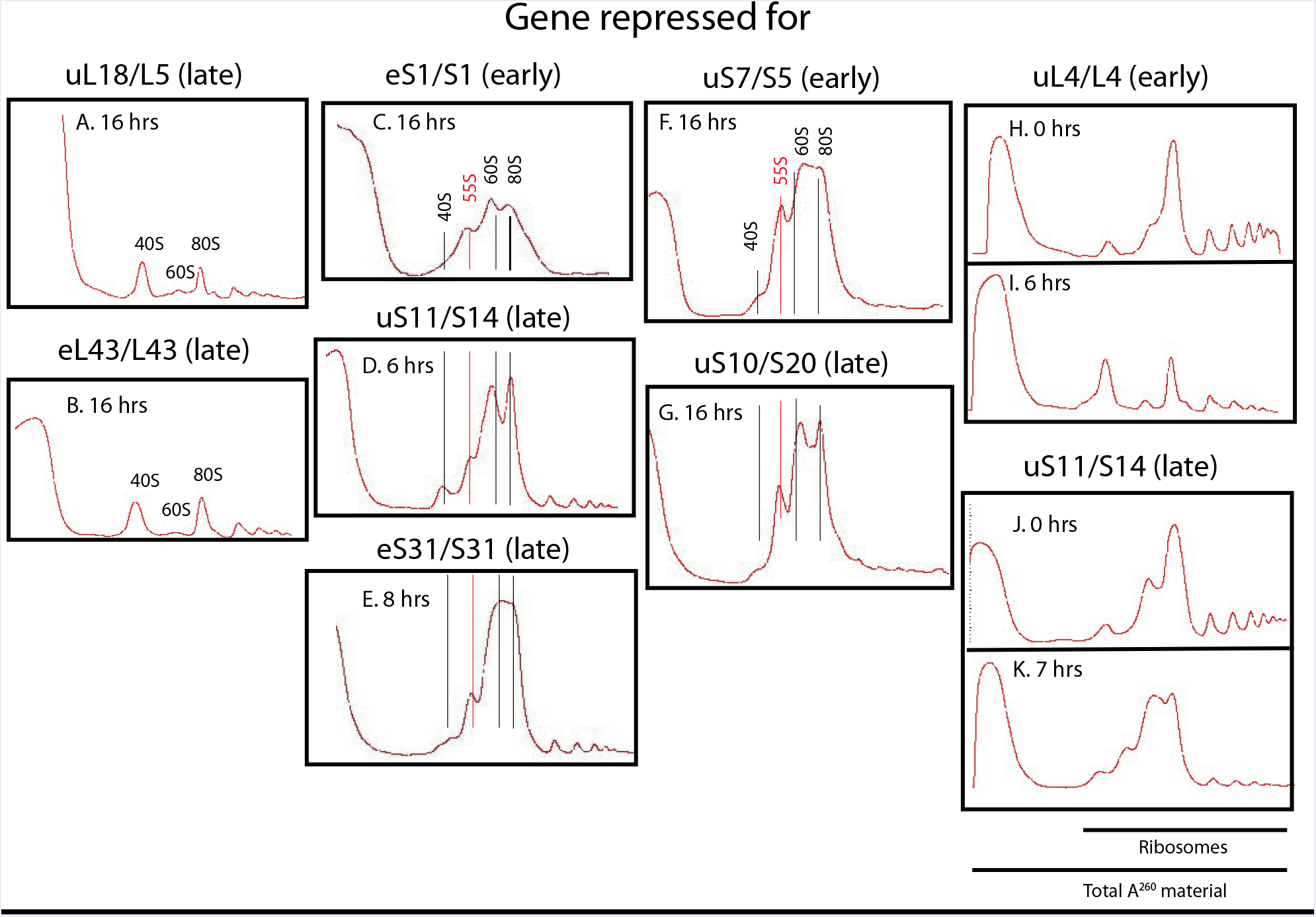
Sucrose gradient analysis of ribosomes before and after cessation of the synthesis of the indicated 60S (A, B, and H-I) or 40S (C-G and J-K) r-proteins. Cells were grown in YEP galactose medium and shifted to YEP glucose medium for the indicated time before harvest. In panels H-L, the spectrophotometer was started a little earlier during gradient tapping to ensure capturing the full peak of A^260^ material at the top of the gradient.

**Supplementary Figure 2.**
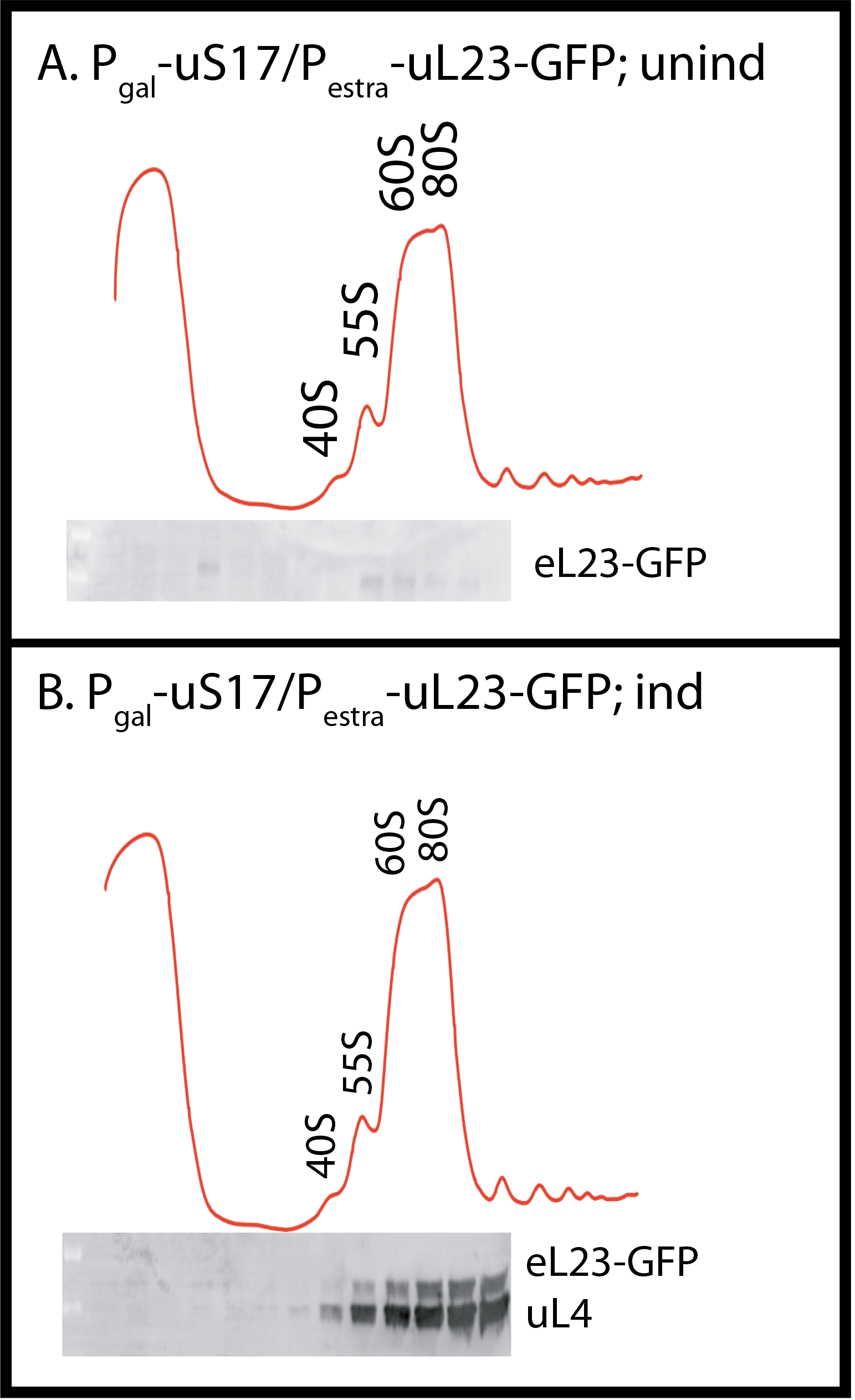
uL23-GFP synthesized during repression of the uS17 gene is incorporated into 80S ribosomes. P_gal_-uS17 carrying a low copy plasmid with an estradiol-inducible uL23-GFP gene was grown in synthetic galactose medium and shifted to synthetic glucose medium. Eight hours after the shift, uL23-GFP synthesis was induced for one hour. Whole cell lysates of cultures harvested before and after the induction were fractionated on sucrose gradients. The indicated fractions were probed by western analysis for GFP and uL4 proteins.

**Supplementary Figure 3.**
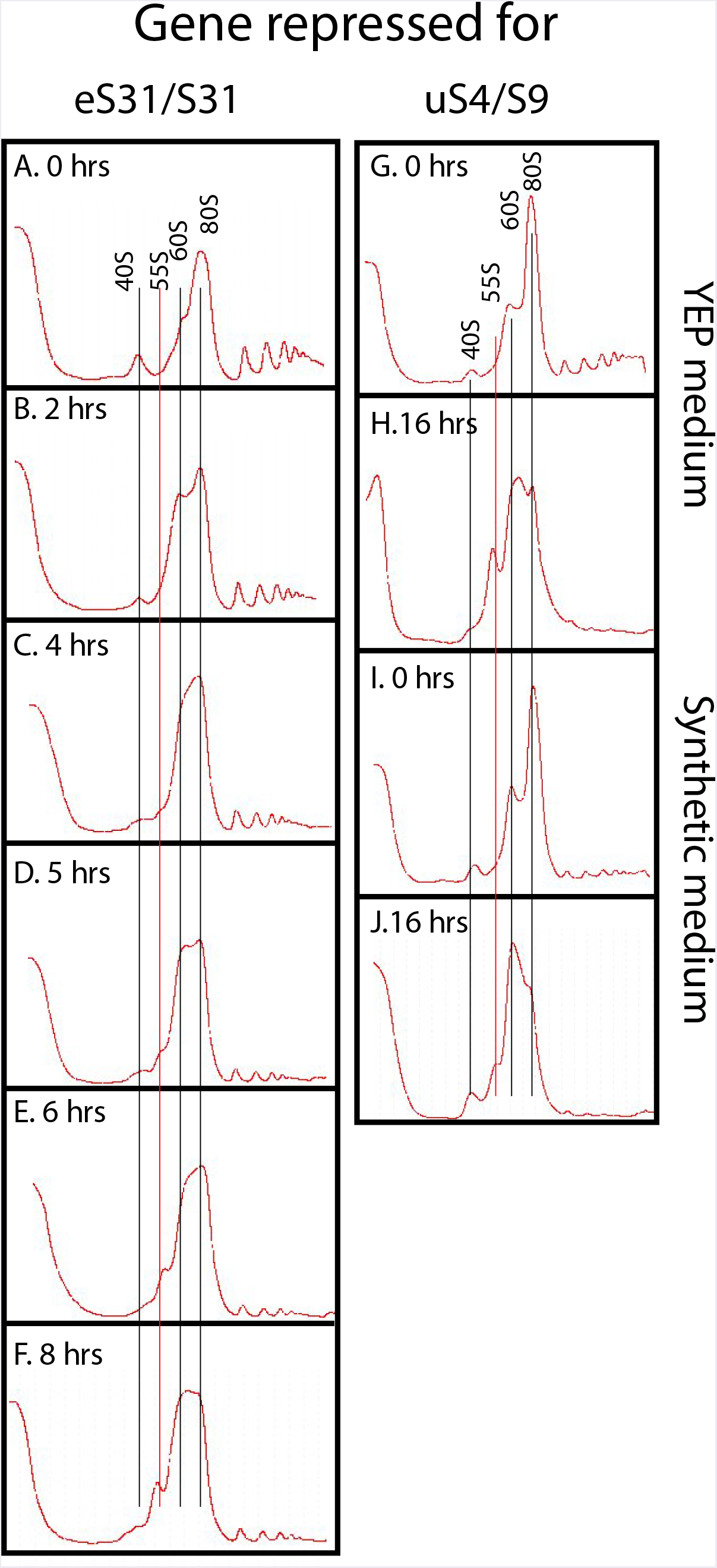
Kinetics of the formation of the 55S peak. A-F. P_gal_-eS31 was grown in YEP galactose medium and shifted to glucose medium for the indicated lengths of time before harvesting. G-J. Effect of growth medium on the level of the 55S particle. P_gal_-uS4 was grown in YEP galactose medium or synthetic complete galactose medium and shifted to the same media, but with glucose as carbon source. G. YEP medium 0 hrs, H. YEP medium 8 hours, I. Synthetic medium 0 hrs, and J. Synthetic medium 8 hours.

**Supplementary Figure 4.**
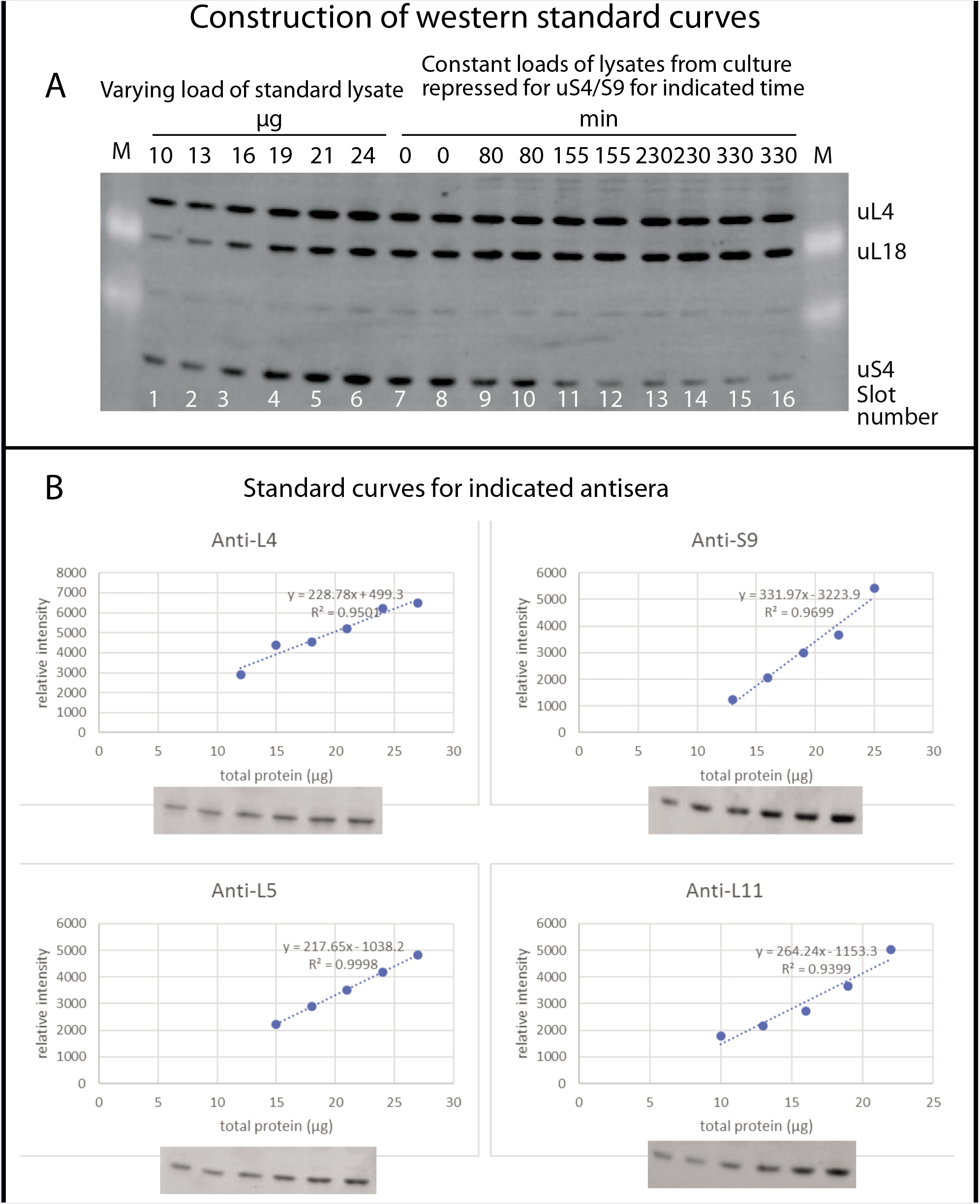
Procedure for quantification of r-proteins by western analysis. A. Slots 1-6 of a 15% SDS polyacrylamide gel were loaded with increasing amounts of whole cell lysate from BY4741 grown in YEP-galactose medium. Slots 7-16 were loaded with constant A^280^ units of whole cell extracts from Pgal-uS4 grown in YEP galactose medium and shifted to glucose medium for the indicated times. After electrophoresis and blotting, the membrane was probed with a cocktail of antisera against uS4/S9, uL4/L4, uL5/L11, or uL18/L5. B. The bands were quantified and the results from slots 1-6 were used to generate a standard curve for each antiserum. The correlation coefficient is indicated for each standard curve. The quantifications of the bands in slots 7-16 were compared to the standard curve and used to quantify the amount of each specific r-protein as a function of time (Figure 4). Note that standard curves were done for each western gel, so that bands in slots 7-16 of each gel could be compared to the standard curve generated from the same blot.

**Supplementary Figure 5.**
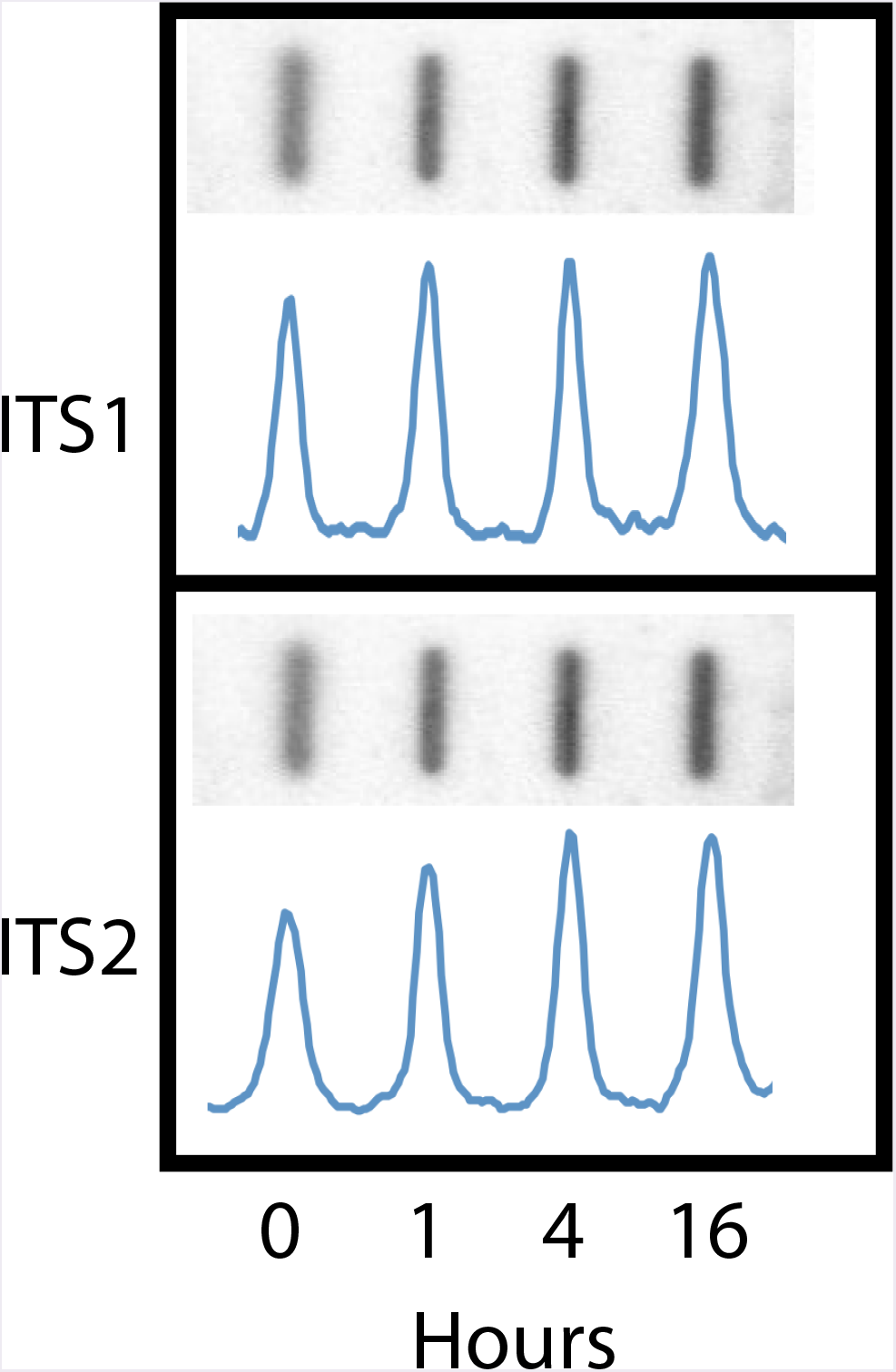
Quantification of rRNA molecules containing ITS1 and ITS2. P_gal_-uS4 was grown in YEP galactose medium and total RNA was prepared from aliquots of culture harvested at 0, 1, 4 and 16 hours after addition of glucose. Equal numbers of A^260^ units of total purified RNA were then loaded onto a nylon membrane using a slot loader. The membrane was probed with ^32^P-labeled oligonucleotides hybridizing to ITS1 (O1663) or ITS2 (O1660); see Figure 3F for specifics about the hybridization probes. The membranes were then scanned on a phosphoimager and the radioactive signal was quantified.

